# The first step is not always the hardest: A change-point analysis of predictive learning

**DOI:** 10.64898/2026.03.17.712476

**Authors:** Nicolas Diekmann, Silke Lissek, Metin Üngör, Sen Cheng

## Abstract

The progress of learning is usually quantified by averaging responses across participants and/or multiple trials within a block. However, such approaches obscure the trial-by-trial progress of learning, which has been shown recently to express a rich variety of dynamics. An alternative approach which does not suffer from this problem is the detection and analysis of points of behavioral change, i.e., change-point analysis. Using change-point analysis, we reanalyzed data from human participants in different predictive learning tasks in which learned contingencies underwent reversal. We find that responses of individual participants were more accurately characterized by behavioral change points than the average learning curve. Importantly, change points significantly shifted to later trials during reversal learning indicating that reversal learning is more difficult than the initial learning. In a computational model based on deep reinforcement learning, we show that the change point shift required the replay of previous experiences, which in turn depends on the hippocampus. This finding is consistent with studies showing that lesions of the hippocampus yield faster reversal learning. In summary, we reaffirm the importance of the analysis of single participant responses, show that phenomenological learning rates are slower during reversal learning, and provide a theoretical account for this difference.

## 1 Introduction

Survival of an individual depends on the successful learning of the appropriate behavior given the environment’s current reward contingencies. Importantly, these reward contingencies are not usually stable over time and might change gradually over time, e.g., a food source might deplete, or abruptly, e.g., when a predator moves into the area. Adjusting a previously learned behavior requires the unlearning of associations (extinction learning) and the learning of new associations (reversal learning). It has been shown that both extinction learning and reversal learning do not simply erase previously learned associations. Previously learned behavior re-emerges with the passage of time (spontaneous recovery), when the individual is re-exposed to the original acquisition context (renewal), or when the individual unexpectedly receives the same reward/punishment (reinstatement) (Bouton, 2004, 2019).

There are clear indications that extinction and reversal learning are different from the initial acquisition. They are known to progress slower compared to acquisition learning (Quirk, 2002), are attenuated by presence (McDonald et al., 2001), number (Bustamante et al., 2016; Wong et al., 2023; Chao and McConnell, 2024) and quality (Wright et al., 2009; Skinner et al., 2014; Wright et al., 2019) of extinction/reversal contexts, and depend on the reinforcement schedule during acquisition training (Amsel, 1967; Shull and Grimes, 2006). However, quantifying learning behavior is a tricky business. A common method is the estimation of learning rates from participants’ responses. To this end, responses are usually averaged across participants when estimating learning rates for a whole group (Üngör and Lachnit, 2006, 2008) or averaged over a sliding window, blocks or sessions when estimating learning rates for single participants (Guo et al., 2014). Averaging data from a few participants does not necessarily yield a good estimate of learning behavior, and measurement noise and outliers will have a large effect on the average. Furthermore, the process of averaging obscures the actual dynamics of learning, which can occur within few trials, and ignores potential intra- and inter-individual differences (Gallistel et al., 2004; Moore and Kuchibhotla, 2022). More recently, studies focusing on the trial-by-trial analysis have revealed rich variability in the learning dynamics of extinction learning. For instance, Donoso et al. (2021) analyzed the trial-by-trial choice behavior of individual pigeons in repeated sessions of an ABA extinction paradigm and found that behavior changed rapidly within sessions and varied significantly between sessions.

The trial-by-trial analysis of behavior also facilitates investigations of the cognitive processes and neural mechanisms underlying extinction and reversal learning. For instance, a recent study of trial-by-trial neural activity associated with reward prediction error signals revealed that these signals occurred during reward omission during early extinction trials and shifted as extinction learning progressed (Packheiser et al., 2021). In another example, a model-driven analysis of human fMRI data, which used trial-by-trial prediction error values derived from a reinforcement learning (RL) model, revealed the involvement of the cerebellum in processing the prediction error (Batsikadze et al., 2022). However, the dynamics of extinction and reversal learning at high temporal resolutions, e.g., trial-by-trial, still remain to be elucidated, and understanding how learning progresses at this level is essential to develop effective interventional methods and the analysis of neuronal recordings.

One particularly interesting neural structure for extinction and reversal learning is the hippocampus, because both are strongly context-dependent (Bouton, 2004; Bouton et al., 2011) and the hippocampus is required for mediating this context-dependence (McDonald et al., 2002; André and Manahan-Vaughan, 2016). While not focusing on the trial-by-trial dynamics, Walther et al. (2021) found that context-dependence spontaneously emerged in an RL model that was not given explicit context information and that this process required replay, a phenomenon closely linked to the hippocampus. When impairing replay in their RL model Walther et al. (2021) they found faster extinction learning and a lack of the context-dependent renewal response.

However, trial-by-trial analyses of learning are difficult to apply when learning is fast, i.e., learning occurs within a small number of trials. A low number of trials, be it in total or for a specific learning condition, limits the window size and effectiveness of the sliding window average. An alternative approach to the quantification of learning behavior is the identification of behavioral change points, i.e., change-point analysis (Gallistel et al., 2004). Using this method, we re-examined human choice responses in predictive learning tasks (Üngör and Lachnit, 2006, 2008; Bustamante et al., 2016; Uengoer et al., 2020), in which learning is too fast to allow for the estimation of individual learning rates. We focused on estimating behavioral change points in acquisition and extinction/reversal learning phases. Our analysis is supplemented with a computational study which gives a theoretical account of the observed learning dynamics and their dependence on the hippocampus.

## 2 Results

### 2.1 Learning in individuals is not gradual but abrupt and switch-like

We re-examined responses from human participants in prediction learning tasks employing similar reversal (Üngör and Lachnit, 2006, 2008; Uengoer et al., 2020) and extinction learning paradigms (Bustamante et al., 2016). Participants were presented different stimuli, i.e., food items, in different contexts, i.e., restaurants, and asked to predict whether a given stimulus would result in stomach trouble. These associations between stimulus/context compounds and outcomes were stable in the acquisition phase and were learned rapidly by participants (Fig. 1). In the reversal learning studies, the outcome associations of either one (Üngör and Lachnit, 2006, 2008) and two (Uengoer et al., 2020) stimulus pairs underwent reversal training, i.e., a stimulus that previously predicted stomach trouble stopped doing so after a context switch and vice versa. In the extinction learning study, one stimulus stopped predicting stomach trouble in the extinction phase (Bustamante et al., 2016). Note that this extinction paradigm nevertheless also requires a change in behavior and is hence comparable to the studies employing a reversal paradigm. During acquisition and reversal/extinction phases, the participants received feedback on the correctness of their responses.

**Figure 1.**
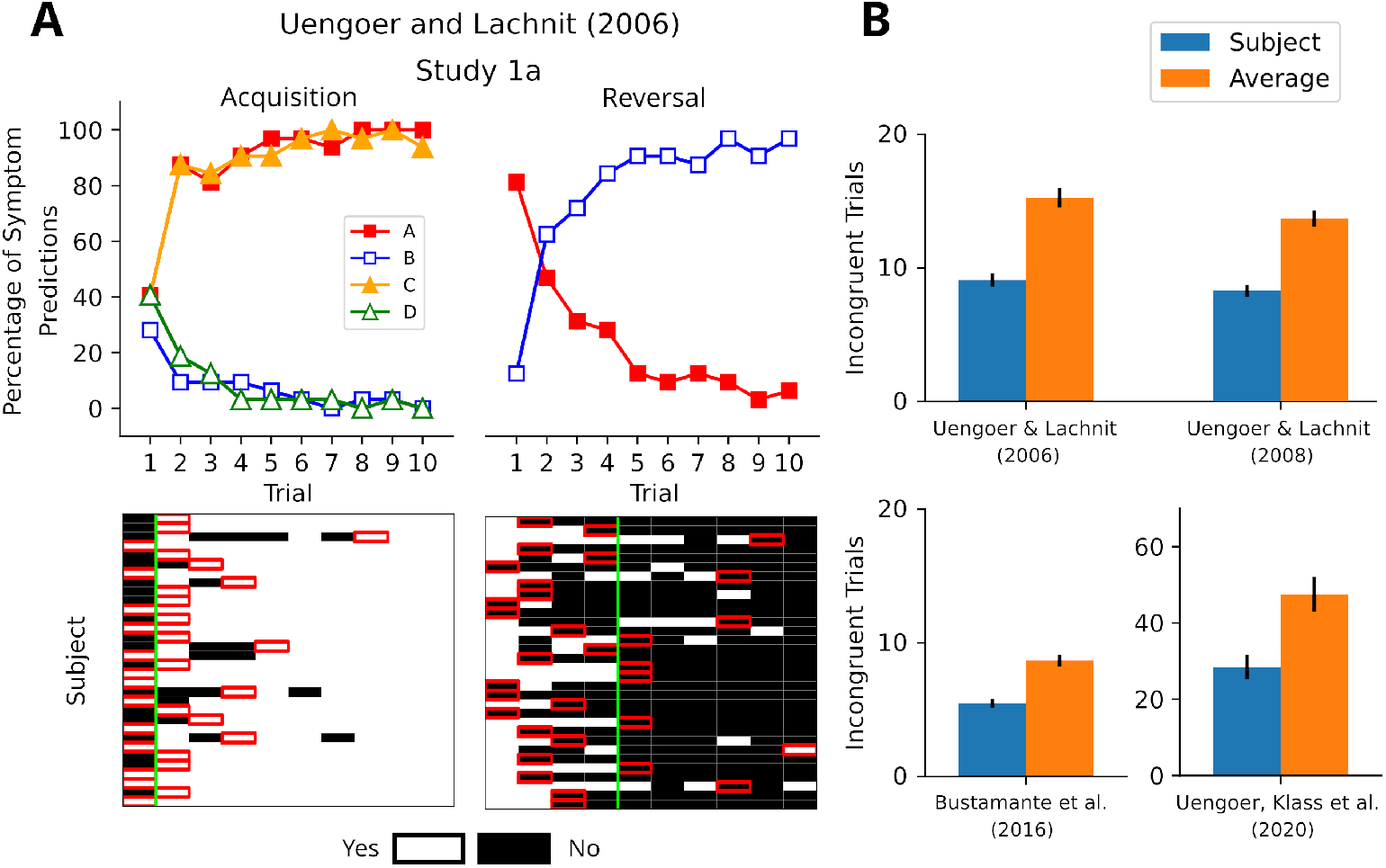
Change-point analysis gives a better account of behavior than average learning curves. **A**: Top: Replotting of Figure 1 from Üngör and Lachnit (2006) (Group Reversal only) which shows the average fraction of positive responses (“yes” responses) for stimuli A-D during phase 1 (acquisition) and phase 2 (reversal). The average responses appear as relatively smooth learning curves. Bottom: The participants’ responses for stimulus A (see Figure S1 for the other stimuli) depicted using a binary coding (white = “yes”, black = “no”). Individual participants appear to change their responses abruptly, not gradually. Identified per-participant change points are marked with red squares, the change points identified based on the average response are marked with a green vertical line. **B**: Single-participant change points produce fewer incongruent trials (correct responses before change point or incorrect responses after) indicating that they better describe the actual behavior for all four experiments.

The original studies averaged the trial-by-trial responses for each stimulus across the participants resulting in apparently smooth learning curves (Fig. 1A, top). However, inspection of the single-participant responses paints a different picture. During the initial acquisition phase, participants predominantly exhibited one of three behaviors: 1. They selected the correct answer on the first trial and did not change it on subsequent trials. 2. They selected the wrong answer on the first trial and switched to the correct one from the second trial onward. 3. They made more than one error before learning the correct answer (Fig. 1A, bottom; S1). During reversal, participants showed similar switching behavior, albeit with more variance between their responses. Importantly, this switching behavior was obscured in the average response curves which convey a false impression of gradual learning. Instead, learning curves are better understood as describing the percentage of participants who have switched their responses.

Quantifying the learning dynamics usually relies upon estimating a learning rate from trial responses. However, the data re-examined here are not conducive to this approach as participants exhibited switching behavior. Therefore, we instead quantified the learning dynamics of single participants by performing a change-point analysis. For each stimulus in the acquisition and reversal/extinction phases (Fig. 1A, bottom), behavioral change points were determined by using binary segmentation (see Methods). Recall phases, if present, were excluded from this analysis due to the lack of response feedback. Behavioral change points detected in the acquisition phase did not differ significantly between the studies (Kruskal-Wallis test, *H* = 5.544, *p* = 0.476). To compare how well the detected change points described the participants’ behavior as compared to the average learning curve, we calculated the number of incongruent trials. Since the change point reflects the trial, in which the contingency was learned, a trial was considered incongruent if a correct response occurred before the change point or an incorrect response after the change point. We found that for all of the re-examined studies, the per-participant behavioral change points produced fewer incongruent trials than the change points based on average learning curves (Fig. 1B). Hence, individual change points offer a more accurate description of the participants’ behavior compared to the averaged responses.

### 2.2 Reversal learning is slower compared to acquisition learning

Previous studies in rodents have shown that learning is slower during reversal in spatial navigation tasks (McDonald et al., 2001) and during extinction learning in a fear conditioning paradigm (Quirk, 2002). Indeed, we were able to find a similar effect for the behavioral change points associated with stimuli undergoing reversal. We performed a type II ANOVA on the change points detected for reversal stimuli from the studies employing the reversal paradigm considering stimulus and phase as factors. We found no significant effect for stimulus (*F* = 1.029, *p* = 0.310) or the interaction stimulus × phase (*F* = 0.019, *p* = 0.890), but a significant effect for phase (*F* = 170.010, *p* = 6.283 × 10^−37^). To assess the differences of change points in acquisition and reversal phases, we performed pair-wise permutation tests for each study. While behavioral change points tended to occur during the first two trials during acquisition they occurred at later trials during reversal (Fig. 2A-C, S2), and this effect was significant and consistent across all re-examined studies (Fig. 2D-F). For non-reversal stimuli such shifts of behavioral change points did not occur.

**Figure 2.**
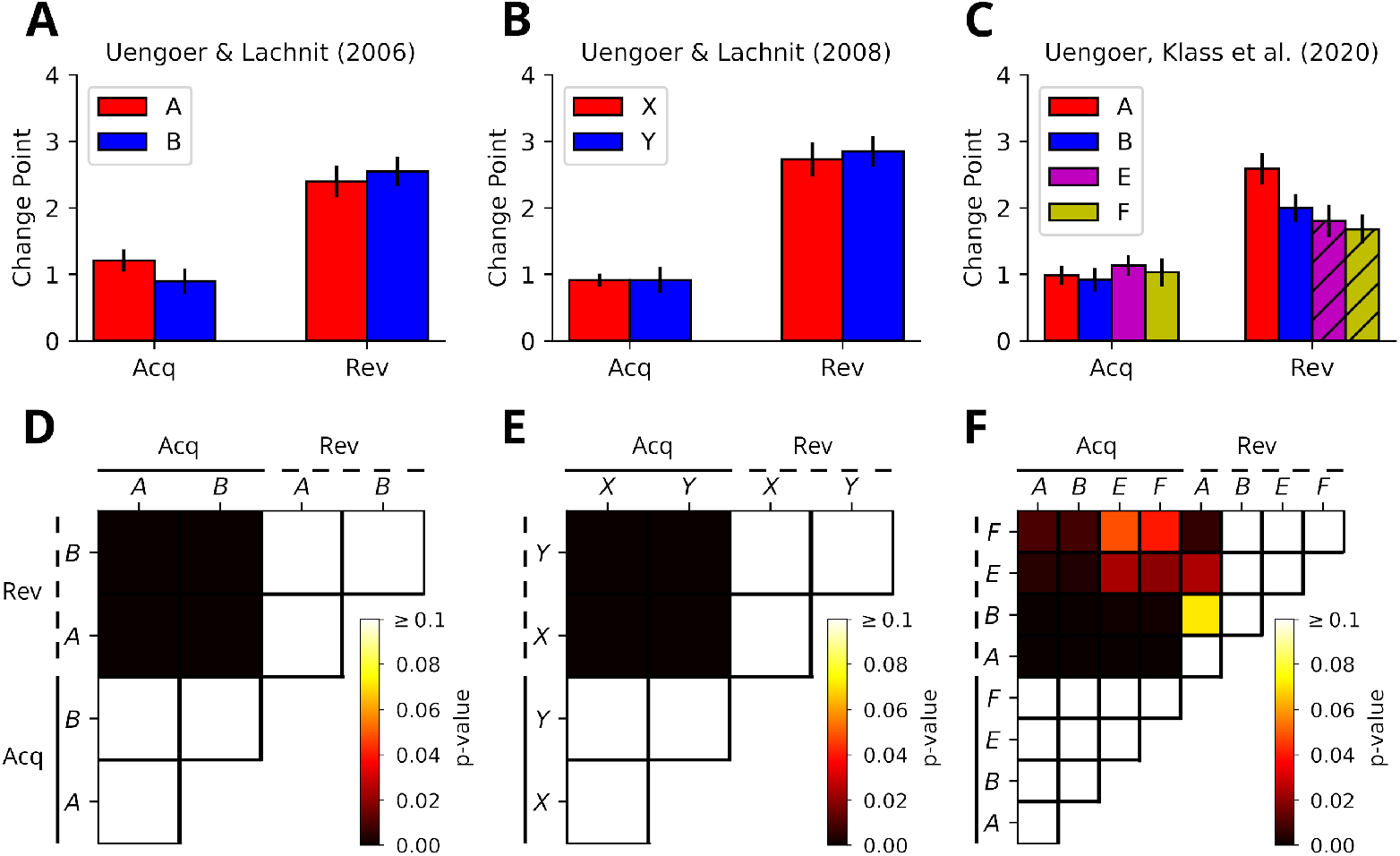
Reversal learning is slower compared to acquisition learning. **A**: The average change points for the reversal stimuli during acquisition (Acq) and reversal (Rev) in Üngör and Lachnit (2006) (groups are shown separately in Fig. S2A). During acquisition the change points tend to occur around the first trial while during reversal they do so after two to three trials. **B**: Same as **A** for data from Üngör and Lachnit (2008) (groups are shown separately in Fig. S2B). **C**: Same as **A** for data from Uengoer et al. (2020). Striped bars indicate that reversal occurred in a different context. **D-F**: Heatmaps depicting the p-values obtained from pair-wise permutation tests of change points for reversal and control stimuli in acquisition and reversal phases for Üngör and Lachnit (2006, 2008); Uengoer et al. (2020), respectively. Generally, change points significantly differ between phases, but not within phases. The only exception is the significant difference between stimulus A and stimuli E-F during reversal in panel F.

Notably, while change point shifts appear to be mostly consistent across studies, the change points detected for reversal stimuli in Uengoer et al. (2020) show some small differences. Firstly, compared to Üngör and Lachnit (2006, 2008) the magnitude of the shift is less pronounced 2C). Secondly, the shift is significantly different between stimulus A and stimuli E and F (Fig. 2F). We may consider three possible explanations for these effects. First, Uengoer et al. (2020) had participants experience multiple sessions of reversal learning, albeit with each session having distinct stimuli, and participants could be expressing faster reversal learning due to meta-learning. A problem with this explanation is that we would expect all stimuli to be affected in the same way. Second, meta-learning or generalization occurs due to two reversal pairs being present in this study’s design. This explanation likewise should predict similar effects on learning speed for all stimuli. Alternatively, the stronger shift for stimulus A could be due to it consistently being the first stimulus presented in the reversal phase. However, we did not find this to be the case (*A*_*first*_ = 21, *B*_*first*_ = 30, *C*_*first*_ = 20, *D*_*first*_ = 17). Third, stimuli A and B undergo reversal without a change of context while stimuli E and F undergo reversal following a change of context. Reversal learning has been shown to be slower when occurring in the same context (McDonald et al., 2001), and therefore the difference between stimuli could be due to whether a context change occurred or not. However, in this case we would expect the shift for stimulus B be comparable to that of A. In summary, the small deviation of Uengoer et al. (2020) remains puzzling.

### 2.3 The effects of context changes on learning speed

Context is known to affect the speed of learning as well as the strength of renewal (Bustamante et al., 2016). Importantly, different environmental features were shown to drive learning to varying degrees (Wright et al., 2009; Skinner et al., 2014; Wright et al., 2019). Of the studies we re-examined that employ reversal learning, it is always accompanied with a context shift except for experimental group AAB from Üngör and Lachnit (2008) and stimuli E and F in Uengoer et al. (2020). To test for potential effects of contexts on reversal learning we pooled detected behavioral change points of the reversal stimuli from Üngör and Lachnit (2006, 2008) and compared same-context and context-change conditions. A one-way ANOVA did not yield a significant difference (*F* = 3.249, *p* = 0.072). Detected behavioral change points from the reversal phase of Uengoer et al. (2020) were excluded from pooling due to the aforementioned differences in the reversal phase. Note that when included the ANOVA yields strongly different result at *p* = 0.149 (*F* = 2.087). While behaviorally change points visually appear to show larger shifts during reversal in the same context as compared to changing contexts, the difference did not reach significance in permutation tests (*p* = 0.081, Fig. 3).

**Figure 3.**
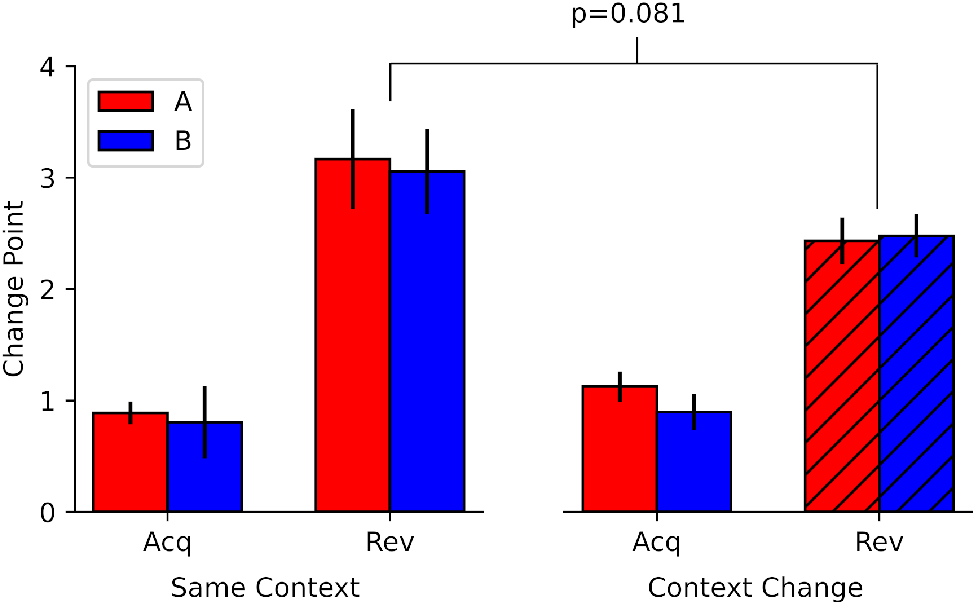
The effects of context changes on reversal learning speed. Change points pooled from Üngör and Lachnit (2006, 2008) for context change and same-context reversal. Striped bars indicate that reversal occurred in a different context. While the change point shift during reversal appears to be larger when reversal occurs in the same context than in different context, a permutation test does not reach a significant threshold (*p* = 0.081).

While the literature suggests that context change leads to fast extinction learning, recent studies show that extinction learning is slower when extinction training occurs in multiple contexts while leading to stronger generalization of extinction (Bustamante et al., 2016, 2024). Reanalyzing the behavioral data from Bustamante et al. (2016) using our change-point analysis yielded a better description of the participants’ behavior and we wondered whether the change-point analysis can reproduce the differences between single- and multiple-context extinction. We pooled detected change points of the extinction stimulus within each extinction schedule and phase, and performed pairwise permutation tests. Indeed, there were significant differences between the change points in the single- and multiple-context extinction schedules as well as significant differences between acquisition and extinction (Fig. 4).

**Figure 4.**
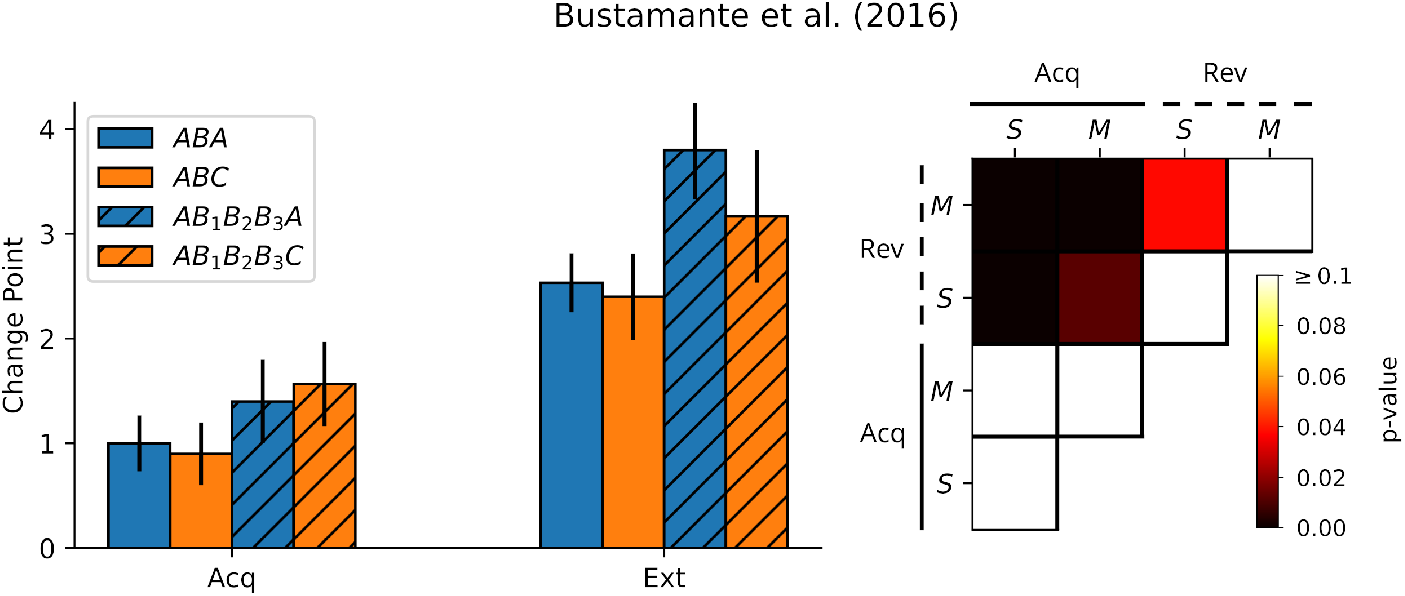
Using multiple contexts slows down extinction learning. **Left**: Average change points for the extinction stimulus during acquisition (Acq) and extinction (Ext) phases from Bustamante et al. (2016). Stripes indicate that extinction was performed using multiple contexts. **Right**: Heatmaps depicting the p-values obtained from permutation tests comparing change points for single- (S) and multiple-context (M) conditions in acquisition and extinction phases. Change points significantly differ between acquisition and extinction, and between single- and multiple-context extinction conditions.

### 2.4 Replay in a reinforcement learning model explains slower learning during reversal

Due to its recent success in accounting for context-dependence of extinction learning (Walther et al., 2021) and replicating human learning dynamics (Batsikadze et al., 2022), we chose the deep-Q network (DQN) (Mnih et al., 2015) to model the potential neural mechanisms underlying the responses of the studies’ participants (Fig. 5). Importantly, we fitted the model’s learning hyper-parameters to the participants’ responses (see Methods) to ensure that it captures the trial-by-trial learning dynamics. Applying the same change-point analysis to the simulated responses of the fitted model we find that they exhibit the same shifts of behavioral change point from acquisition to reversal phase (Fig. 6, left).

**Figure 5.**
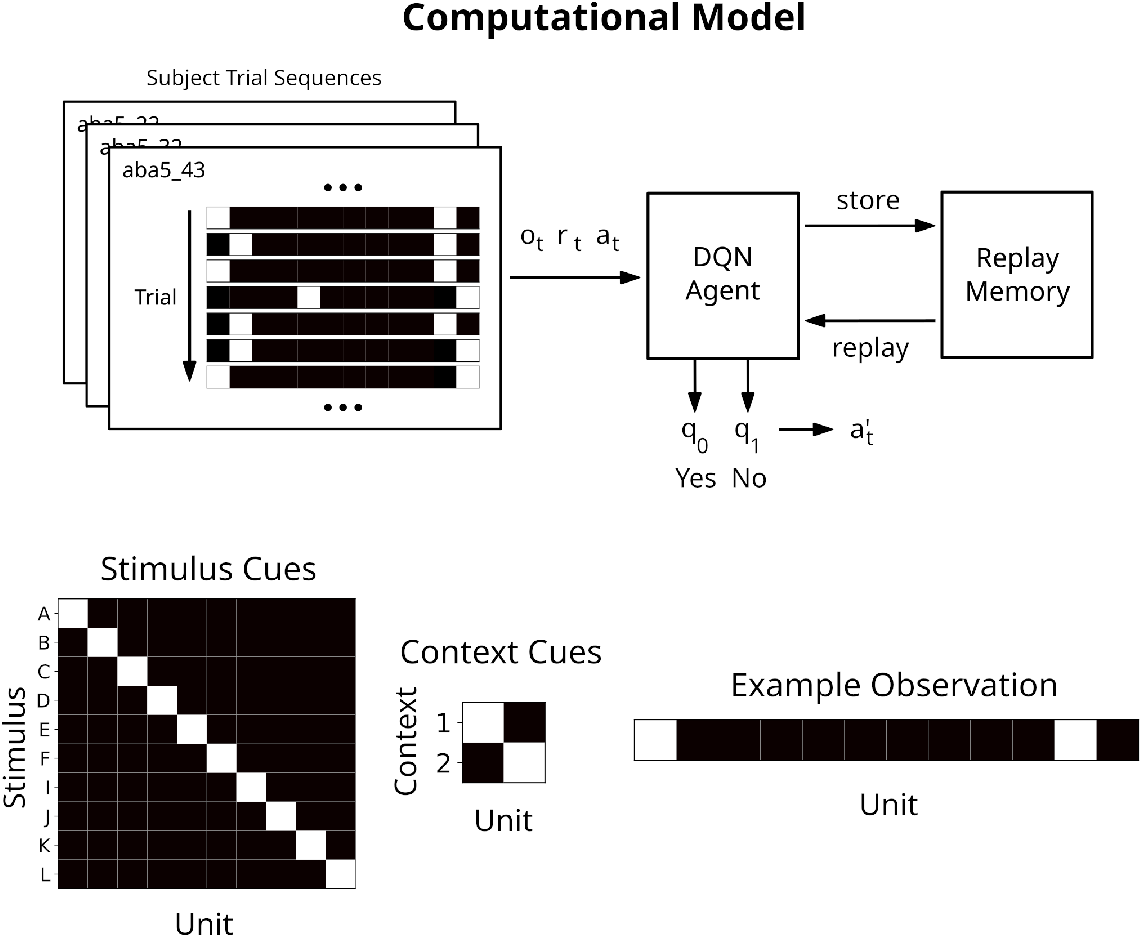
Illustration of the computational model. **Top**: The RL simulation setup. Unique trial sequences for each participant are presented to the DQN agent. The DQN agent makes predictions for actions on each trial 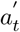, but executes the actions *a*_*t*_ dictated by the trial sequences. **Bottom**: Example observations for study 1a from Üngör and Lachnit (2006). Stimuli and contexts are represented using a one-hot encoding. Their representations are concatenated and fed as input to the network. The example input on the right corresponds to stimulus A in context 1.

**Figure 6.**
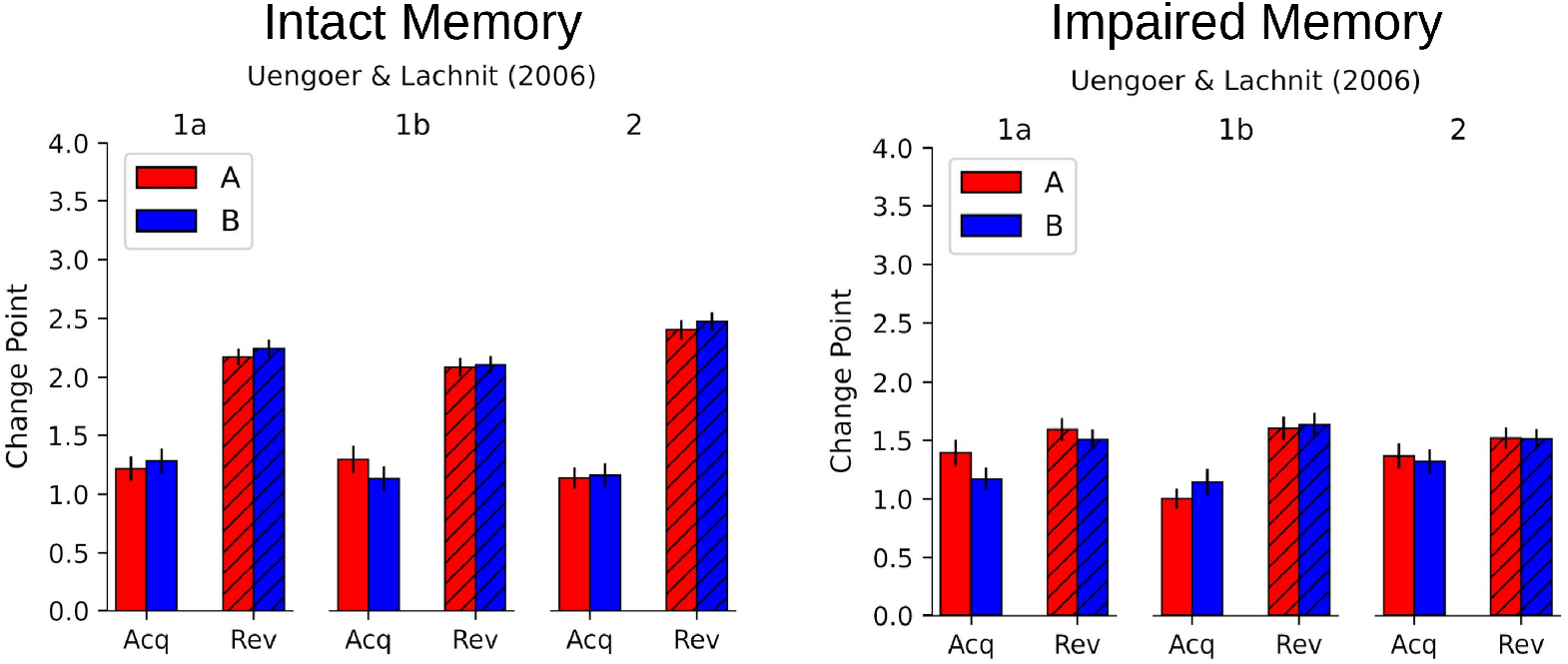
Replay in a reinforcement learning model accounts for slower learning during reversal. **Left**: Behavioral change points obtained from simulations based on the model that best-fit the data from Üngör and Lachnit (2006). Behavioral change points in the model shift during reversal just as in the experimental data. **Right**: If the model’s replay memory is impaired, behavioral change points do not shift during reversal.

The speed of both extinction and reversal learning are known to be attenuated in the presence of an intact hippocampus (McDonald et al., 2002). Walther et al. (2021) used replay-driven deep RL to model the role of hippocampus in context-dependent extinction learning, and argued that the hippocampus facilitates context-dependent learning by replaying experiences from acquisition and extinction contexts. Intriguingly, their model also shows strong attenuation of extinction learning speed due to replay of older experiences, and impairment of replay memory resulted in faster extinction learning and vanishing of the renewal effect. Limiting the replay memory in our model to only those experiences from the current learning phase (impaired memory), we found that behavioral change points stop shifting during reversal (Fig. 6, right). These results suggest that the hippocampus might also modulate learning rates in different experimental phase throught its role in driving replay.

As we mentioned above, in our hands there is trend-level evidence that reversal learning in the same context in slower than in different contexts (Fig. 3). More definitively, studies in rats show that the similarity of context modulates the speed of reversal learning (McDonald et al., 2001; Bouton et al., 2011). However, such an effect was not detectable in our simulations using the learning hyper-parameters obtained from our grid search. We hypothesize that this is because learning was too fast and the prioritization of recent trials over past ones removes interference due to context similarity. This recency prioritization was introduced into a previous model (Batsikadze et al., 2022) for this purpose and replaced a more artificial mechanism in an antecedent model, where the replay memory capacity was somewhat arbitrarily limited. In both cases, the purpose was to accelerate the re-extinction of the renewed response, which otherwise suffered from interference by similar experiences from the acquisition phase (Walther et al., 2021). This, however, is precisely what we need to account for the subtle difference between reversal learning in the same vs. different context. Slowing down learning in the model by adapting the learning hyper-parameters, we found that the absence of a context change indeed attenuates learning speed (Fig. S6).

To better understand how replay affects learning in the network, we recorded weight changes in the network as well as predicted Q-values for the stimuli in both acquisition and reversal contexts. Our simulations show that network weights show similar network changes throughout the acquisition phase for agents with intact, i.e., experiences from all phases can be replayed, and impaired memory (Fig. 7A). In contrast during reversal, networks with intact memory show an initial spike in network changes followed by a prolonged phase of smaller changes whereas networks with impaired memory show only one large spike of network changes early in the reversal phase (Fig. 7A). Recorded Q-values show correct learning of the reward contingencies and their generalization to the reversal context during the acquisition phase in both settings (Fig. 7B). In the reversal phase, the RL agent with intact memory showed that parallel to the slow learning of the new contingencies, the previously learned contingencies for the acquisition context underwent a prolonged phase of de- and re-stabilization. The RL agent with impaired memory did not show any of this and, instead, quickly learned the new reward contingencies in the reversal context and forgot the previously learned associations, suggesting that catastrophic interference occurred between the two sets of contingencies. Based on these and previous (Walther et al., 2021; Batsikadze et al., 2022) simulation results, we argue that slower learning speeds during reversal can be explained by interference between experiences from acquisition and reversal phases, which are replayed with the help of the hippocampus. This interference induces the necessary synaptic changes to represent contextual information and is accompanied by a brief phase of destabilization of existing memories.

**Figure 7.**
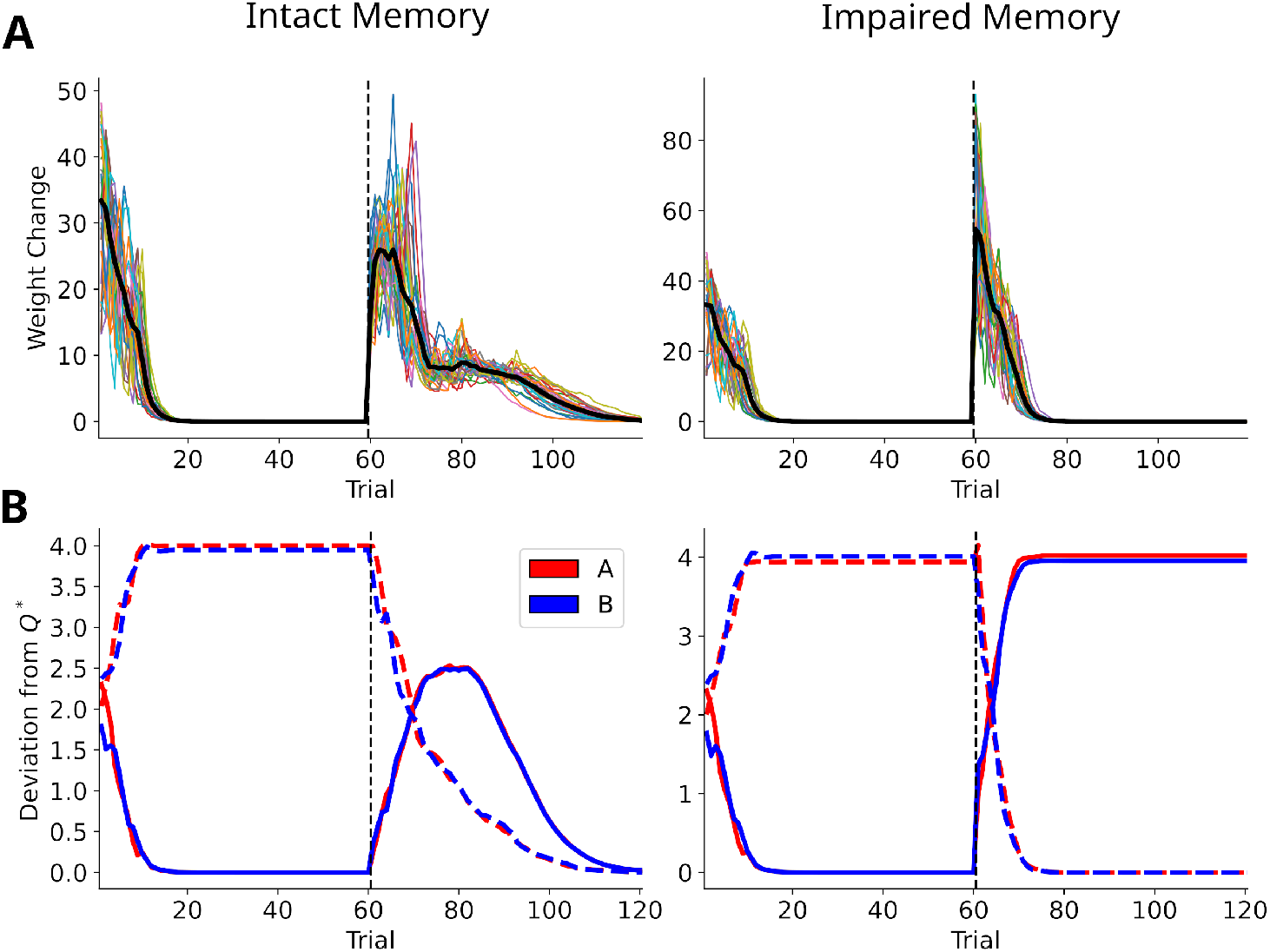
Mechanism underlying shift in change point during reversal learning. **A**: The trial-to-trial network weight changes for RL agents with intact (left) and impaired (right) memory. During acquisition, the networks undergo similar changes, while during reversal the intact memory agent shows a more extended period of medium and small weight changes. **B**: Deviation from the optimal Q-values (*Q*^∗^) for the reversal stimuli in both contexts over the course of learning with solid and dashed lines representing acquisition and reversal context, respectively. The network initially learns the reward contingencies in the acquisition phase and generalizes them to the reversal context. The intact memory agent (left) learns the new context-dependent contingencies in the reversal phase during which the Q-values for the stimuli in the acquisition context become destabilized. In contrast, the impaired memory agent unlearns the acquisition context contingency as it learns the reversal context contingency.

## 3 Discussion

We identified behavioral change points on a single-participant level in human behavioral data of four studies employing similar predictive learning tasks. In agreement with recent critiques of the average learning curve, we found that individual change points better described the behavioral data than the average learning curve. Importantly, we found that behavioral switches occurred later during the reversal phase compared to the acquisition phase. Finally, we modeled the experimental data with a deep reinforcement learning (RL) model, which was driven by experience replay and used the same underlying learning rates for both acquisition and reversal phases. Our model replicated the shifts of behavioral change points during reversal and demonstrated that experience replay is key for this effect.

### 3.1 The average learning curve and gradual learning

The approach of averaging responses across individuals and/or sessions/trials has been criticized by several authors (Gallistel et al., 2004; Glautier, 2013; Donoso et al., 2021). Most prominently, Gallistel et al. (2004) argued that the graduality of learning is but an artifact of the averaging process, and, hence, average learning curves cannot be a reliable measure of learning rate. Among their re-analysis of experimental data they employed the detection of putative change points similar to our analysis. However, there are also contrary views defending the use of gradual learning curves to describe learning in individuals (Harris, 2022; Moore and Kuchibhotla, 2022). For instance, a reanalysis of behavioral data from rats in an appetitive extinction learning paradigm found that acquisition and extinction were best described by gradual decelerated learning rule (Harris, 2022). Alternatively, Moore and Kuchibhotla (2022) argue that instrumental learning can be divided into two parallel processes: a fast step-like acquisition of knowledge and a slow variable expression of behavior.

The experimental results presented here favor Gallistel et al. (2004)’s view since participants showed switch-like behavior, but are also consistent with Moore and Kuchibhotla (2022)’s view, if we make a few assumptions. Namely, the step-like acquisition took place within one trial during initial learning and expression of behavior was delayed by one trial, and later behavioral switches during reversal were due to slower step-like acquisition of the changed contingencies. However, we cannot rule out gradual learning. Given a high enough learning rate (*α* = 1 in the most extreme case) even learning models which are updated using gradual learning rules would produce switch-like behavior. Furthermore, even if behavior has changed, learning may still occur. Consider the simple example of a Rescorla-Wagner model that learns value associations based on which binary choice is made depending on whether the learned value passes a specified threshold. While choices will change as soon as the threshold is passed, learning will still proceed until the correct value is predicted. Indeed, our computational model uses a gradual learning rule and when we analyzed the evolution of network weights and predictions of our model, we found that both keep changing even after choice behavior has already switched. This was especially the case during reversal learning when the correct contingencies were learned late in the phase and acquisition contingencies experience destabilization. Hence our modeling results offer an additional third possibility that is contrary to the view proposed by Moore and Kuchibhotla (2022): Learning is gradual and proceeds throughout, and step-like changes occur only at an behavioral level.

In summary, abrupt changes in behavior can emerge from gradual learning rules. However, the gradual average learning curves should generally not be interpreted as a measure of the underlying learning process since learning is most likely subject to intra- and inter-individual differences. In our view, average learning curves are best viewed as representing the fraction of participants that have already switched their behavior at different timepoints of the learning process.

### 3.2 The effect of context on reversal and extinction learning speed

Our re-analysis did not find a significant effect of context on reversal learning. A potential reason for this is that the sample sizes, especially for the change points detected in same-context reversal, were too low, which is further compounded by the fast learning speed. Our results contrast with results reported in rats which show slower reversal learning in the same context (McDonald et al., 2001; Bouton et al., 2011) and different reversal learning speeds depending on the quality of context change (Wright et al., 2009; Skinner et al., 2014; Wright et al., 2019). The lack of a context effect might also be explained by the use of different learning strategies either due to the experimental paradigm, e.g., spatial navigation vs. predictive learning, or species differences, or both. It is noteworthy that a recent study of extinction learning in humans using an operant conditioning paradigm in which one group underwent extinction in the same while another underwent extinction in a different context also did not detect differences in the speed of extinction between the groups (Ritchey et al., 2021). Furthermore, previous studies in rats by Bouton and King (1983) and Bouton and Peck (1989) also did not report an effect of context.

Assuming context did affect learning, but we simply failed to detect it, how could we improve the likelihood of detection? Similar to Ritchey et al. (2021), our analysis included a relatively small sample size for the group undergoing reversal in the same context. We could collect further data using the AAB experimental design from Üngör and Lachnit (2008) and variants of the experiments from Üngör and Lachnit (2006) without context change. Another approach would be adapting or changing the experimental design by making learning overall slower, thereby making the detection of context-dependent differences easier. For instance, the spatial navigation tasks used by McDonald et al. (2001) could be adapted for human participants, e.g., using a virtual reality setup. Task difficulty could also be increased by making the outcomes probabilistic. A potential confound would be that both partial reinforcement and variable reinforcement rate decrease the speed of extinction learning.

A better experimental design for the detection of context-dependent differences could be achieved using a model-driven approach. For instance, the model in Batsikadze et al. (2022) was used to derive a new experimental design that was optimized to increase the magnitude and occurrence of reward prediction errors signals in a fear extinction paradigm (Doubliez et al., 2025; Nio et al., 2025). A similar approach lead to an experimental design that reduces learning speed, thereby increasing the likelihood of detecting effects of contexts on extinction and reversal learning.

### 3.3 The role of hippocampus in phase-dependent learning speed

The hippocampus is necessary for context-dependent extinction and reversal learning (Bouton, 2004, 2019). While the conventional view is that the hippocampus facilitates context-dependence by inferring and/or representing context (Heald et al., 2023), recent work argues that context is learned via the hippocampus replaying experiences from acquisition and extinction contexts to train neocortex (Walther et al., 2021; Batsikadze et al., 2022). In this novel view, hippocampal replay prevents the catastrophic interference of the learned reward contingencies for the acquisition context. Indeed, the replay of acquisition and reversal experiences in our model induced mild interference in the network throughout the reversal phase and temporally destabilized the learned acquisition contingencies, before restabilizing them. It was this mild and prolonged interference that resulted in delayed behavioral switches during reversal. This delay vanished, when we impaired memory replay, but so did the restabilization of previously learned acquisition contingencies, resulting in catastrophic interference.

A previous model, which also incorporates memory (Glautier, 2013), extended the standard Rescorla-Wagner model with memory buffers that store the previous trial, i.e., stimulus and reinforcement information, and configural cues. Furthermore, the update rule is adapted with a weight matrix that differs for each buffer. Through these modification this model can account for the behavior of individual participants better than a standard Rescorla-Wagner model can. Glautier (2013) does not discuss a biological implementation of his model, but one could argue that the model’s memory buffers could reflect the function of the hippocampus, similar to how experience replay in RL is thought to reflect the function of hippocampal replay (Mattar and Daw, 2018; Khamassi and Girard, 2020; Diekmann and Cheng, 2023). However, the earlier model is more rigid and requires and increasing number of memory buffers unlike the RL model used in this study. Furthermore, it is not clear whether Glautier (2013)’s model account for learning dynamics observed in reversal and extinction learning paradigms since his study only modeled a single learning stage/phase.

### 3.4 Conclusion

Our re-analysis of human behavioral data provides further support for recent critiques of representing learning process with an average learning curve, and the delayed behavioral switches we found during reversal fit well in the literature on extinction and reversal learning. Furthermore, we showed that these inter-phase differences can be accounted for by interference of replayed experiences and do not require phase-specific learning rates.

## 4 Methods and materials

We implemented our analysis code as well as our computational model in the *Python* programming language (van Rossum and de Boer, 1991). Change points were identified using the *ruptures* package (Truong et al., 2020), and statistical analyses were performed using the *SciPy* (Virtanen et al., 2020) *and statsmodels* (Seabold and Perktold, 2010) packages. Reinforcement learning (RL) simulations were implemented using the *CoBeL-RL* framework (Walther et al., 2021; Diekmann et al., 2023), and *PyTorch* (Paszke et al., 2019) was used to implement and train the RL agent’s deep neural networks. All code is available at https://github.com/sencheng/change-point-analysis.

### 4.1 Change-point analysis

To identify behavioral change points, we performed binary segmentation on responses for each stimulus in acquisition and reversal/extinction phases. We define the behavioral change point of participant *i* for a stimulus *x* in phase *p* of session *s* as 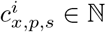. Since only Uengoer et al. (2020) used an experimental design with multiple sessions, we will omit the session subscript *s* for the other studies. For cases in which a participants gave the same response throughout the whole phase for a given stimulus we consider the change point to occur on the initial trial, i.e., 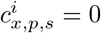. Recall phases were excluded from the analysis as they were too short and participants did not receive any feedback. Additionally, we computed behavioral change points 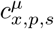 based on the responses averaged across participants. We then computed the number of incongruent responses between the actual responses *a*_*t*_ and synthetic responses 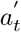 derived from the detected change points.

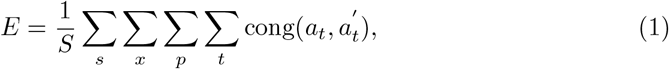

where the function cong measures the congruency of the two trial responses *a* and *a*^′^:

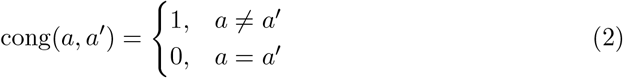

Normality of the distribution of behavioral change points *c*_*x,p*_ was assessed with the test of normality based on D’Agostino and Pearson (1973). The Kruskal-Wallis test was used as an non-parametric alternative to ANOVA since for most *c*_*x,p*_ normality was not given. However, to assess potential interaction effects we relied on two-way ANOVA tests. To identify for which behavioral change points differences occurred, we performed pair-wise two-sided permutation tests based on 100, 000 random reshuffles. We report p-values uncorrected for multiple comparisons in the main text. While correction for multiple comparisons reduces Type I errors it also reduces statistical power, i.e., it increase Type II errors. When we correct p-values using the Holm-Bonferroni method (figures S3-5) we fail to detect significant differences for Uengoer et al. (2020) (figure S4F) and Bustamante et al. (2016) (figure S3). Bustamante et al. (2016) originally reported a significant difference between single and multiple context extinction paradigms, and the reduced power due to corrections is likely the cause for us failing to detect the difference between the two paradigms. Detected change points for Uengoer et al. (2020) shifted less during reversal compared to Üngör and Lachnit (2006, 2008) (except for stimulus A), and the study was therefore likely more affected by the reduction in statistical power.

### 4.2 Computational modeling

We chose RL (Sutton and Barto, 2018) to model the learning dynamics of the participants of the different studies. RL describes the interaction of a reward-maximizing agent with its environment. At each time point *t*, the agent finds itself in an environmental state *s*_*t*_ and performs an action *a*_*t*_ that transitions the agent to a new state *s*_*t*+1_ and yields a real-valued reward *r*_*t*_. We can define each of these interactions as an experience tuple *e*_*t*_ = (*s*_*t*_, *a*_*t*_, *r*_*t*_*s*_*t*+1_). Given these experiences, the agent tries to maximize the expected cumulative reward *R*:

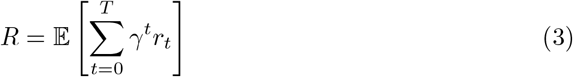

Here, *γ* is the so-called discount factor which weighs the importance of immediate over future rewards. In our simulations, each trial consists only of one step/transition and therefore the *γ* learning hyper-parameter can be ignored.

A popular RL algorithm is Q-learning (Watkins, 1989), which solves the RL optimization problem by learning the expected cumulative reward *Q*(*s, a*), if selecting action *a* in state *s*. At each time step *t*, this Q-function is updated based the current experience *e*_*t*_ and bootstrapping the value of state *s*_*t*+1_ from the current Q-function estimate:

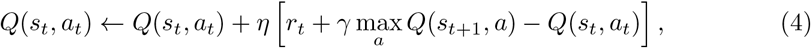

where *η* denotes the learning rate. The term in square brackets is commonly known as the temporal difference (TD) error. Note that for experiences which end in a terminal state, meaning that the trial ended, the bootstrapped value is set to zero.

The original Q-learning algorithm was developed with tabular RL in mind (Watkins, 1989) and is not well suited for modeling context-dependent learning settings and inter-stimulus learning effects. Hence, we used the deep-Q network (DQN) which combines deep neural networks with Q-learning (Mnih et al., 2015), and has been successfully used to model context-dependent extinction learning (Walther et al., 2021; Batsikadze et al., 2022). The DQN represents the Q-function using a deep neural network with parameters *θ*, which is trained via backpropagation (Rumelhart et al., 1986) from experience batches obtained through experience replay (Lin, 1992). For all simulations, the DQN consisted of two hidden fully-connected layers and one output layer. Both hidden layers had 64 units and used the hyperbolic tangent as their activation function. The output layer used two units to convey the two possible actions. Actions were selected following an *ϵ*-greedy action selection policy, that is, the agent selects the action associated with the largest Q-value most of the time, but will select a random action with a small probability *ϵ* (Sutton and Barto, 2018).

The speed of learning for novel information in the DQN partly depends on the number of stored experiences, because experiences are replayed randomly. To allow our model to accurately replicate the learning dynamics of participants we prioritized experiences during replay to favor recent experiences similar to previous work by Batsikadze et al. (2022). The priority rating p_i_ of each experience e_i_ we define as:

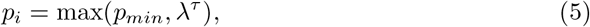

where *λ* is the priority decay factor, *τ* is the time passed in trials and *p*_*min*_ denotes the minimum priority value. The minimum priority value ensures that experiences from the acquisition phase have a reasonable chance to be replayed even for low values of *λ*.

Replay probabilities of experiences *e*_*i*_ are then computed as:

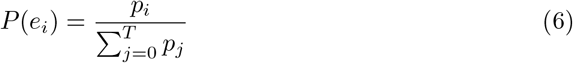

Furthermore, we slightly adapted the update process by providing the network with target values for both actions. This was necessary as some participants would never experience certain state-action combinations which caused the DQN to not update the Q-function for them.

We subjected RL agents to simplified virtual versions of the various predictive learning tasks in which we present stimulus and context as one-hot encoded vectors (Fig. 5). Different instances of an RL agent, that is, instances which share the same learning hyper-parameters, are presented the same unique trial sequences that the participants of the different studies experienced. The RL agents can select either of two actions, i.e., *a*_0_ and *a*_1_, which represent “yes” and “no” answers, respectively. Correct answers yield a positive reward of +1 while incorrect answers yield a negative reward −1. Importantly, while the RL agents form predictions for which action to select *a*_*t*_, the actual actions executed are those originally chosen by the participants, denoted as 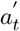.

The learning dynamics of our RL model depends on the different learning hyper-parameters such as the number of replays per trial. We therefore performed a limited grid search over those learning hyper-parameters which most strongly affect the learning dynamics (Table 1): the replay decay parameters *λ*, the batch size *b*, the number of replays *i*, the amount of exploration *ϵ*, the minimum value for experience priority *p*_*min*_ and the initial Q-function *Q*_0_. Each combinations of parameter values and trial sequence was simulated 25 times. We defined the quality of the fit as the average negative log-likelihood over all participants *S*:

**Table 1.**
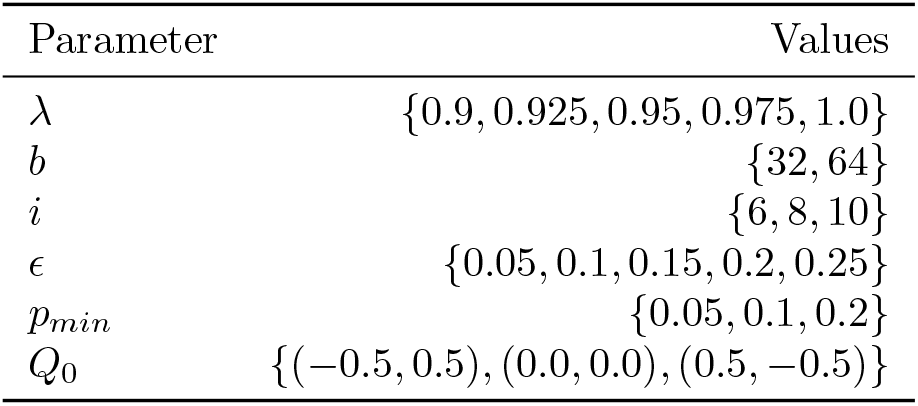
The hyper-parameter values that were part of the grid search. Note that the values for the initial Q-function *Q*_0_ represent the actions in order, i.e., (*a*_0_, *a*_1_).

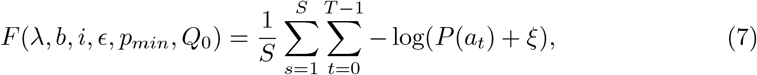

where *P* (*a*_*t*_) is derived from the average of 25 simulation runs. Due to *P* (*a*_*t*_) reaching zero in some cases a small value *ξ* = 10^−^8 is added to the probabilities. The RL model which fitted the participants’ behavior best was then analyzed with the same change-point analysis described above for he experimenal data. We determined the validity of the model based on its ability to replicate results of the change-point analysis of the experimental data, as well as the presence of the renewal effect when applicable.

We performed an additional network analysis in which we compared the evolution of the Q-function and network weight changes for our model with and without replay. Network weight changes Δ*θ* were tracked by computing sum of absolute difference between subsequent trials:

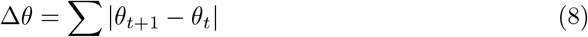

For the evolution of the Q-function we tracked the Q-value prediction for the reversal stimuli at each trial in both contexts.

To test whether our model can express different reversal learning speeds we ran simulations of the studies from Uengoer and Lachnit (2006) with and without context change. The learning dynamics which resulted from the learning hyper-parameters obtained via our grid search were too fast to express a detectable difference. This fast learning was mainly facilitated by the decay factor that was used for replay prioritization. Hence, we adjusted the learning parameters to overall slow down learning.

Specifically, we used a decay factor *λ* = 0.995, a batch size *b* = 64, a number of replays *i* = 10, an exploration rate *ϵ* = 0.15, a minimum priority *p*_*min*_ = 0.05, and initial Q-values *Q*_0_ = (0.0, 0.0).

## 5 Acknowledgments

This study was funded by the Deutsche Forschungsgemeinschaft (DFG, German Research Foundation) — project number 316803389 — SFB 1280, project F01 (SC,MU).

## Supplementary Figures

**Figure S1:**
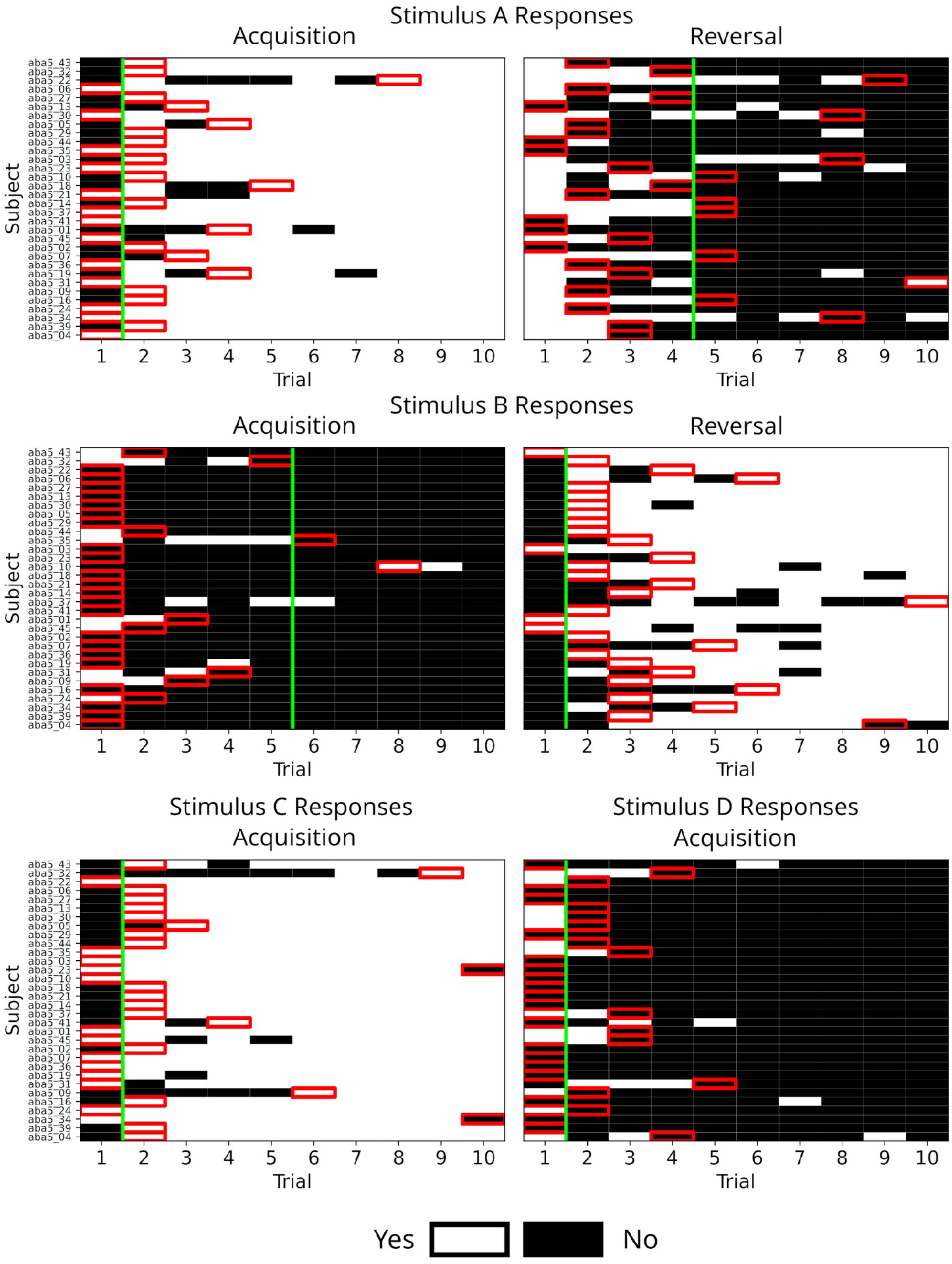
Detailed per-subject responses and change points. Shown are per-subject responses for stimuli A-D in the reversal group of Study 1a from Uengoer and Lachnit (2006). Stimuli A and B served as the reversal pair and were shown during acquisition and reversal phases while stimuli C and D served as control stimuli and were not shown during reversal. Per-subject change points were detected for each stimulus in each phase together with a change point based on the averaged responses. The detected change points are marked with red rectangles, the change point based on the averaged responses in marked with a green line.

**Figure S2:**
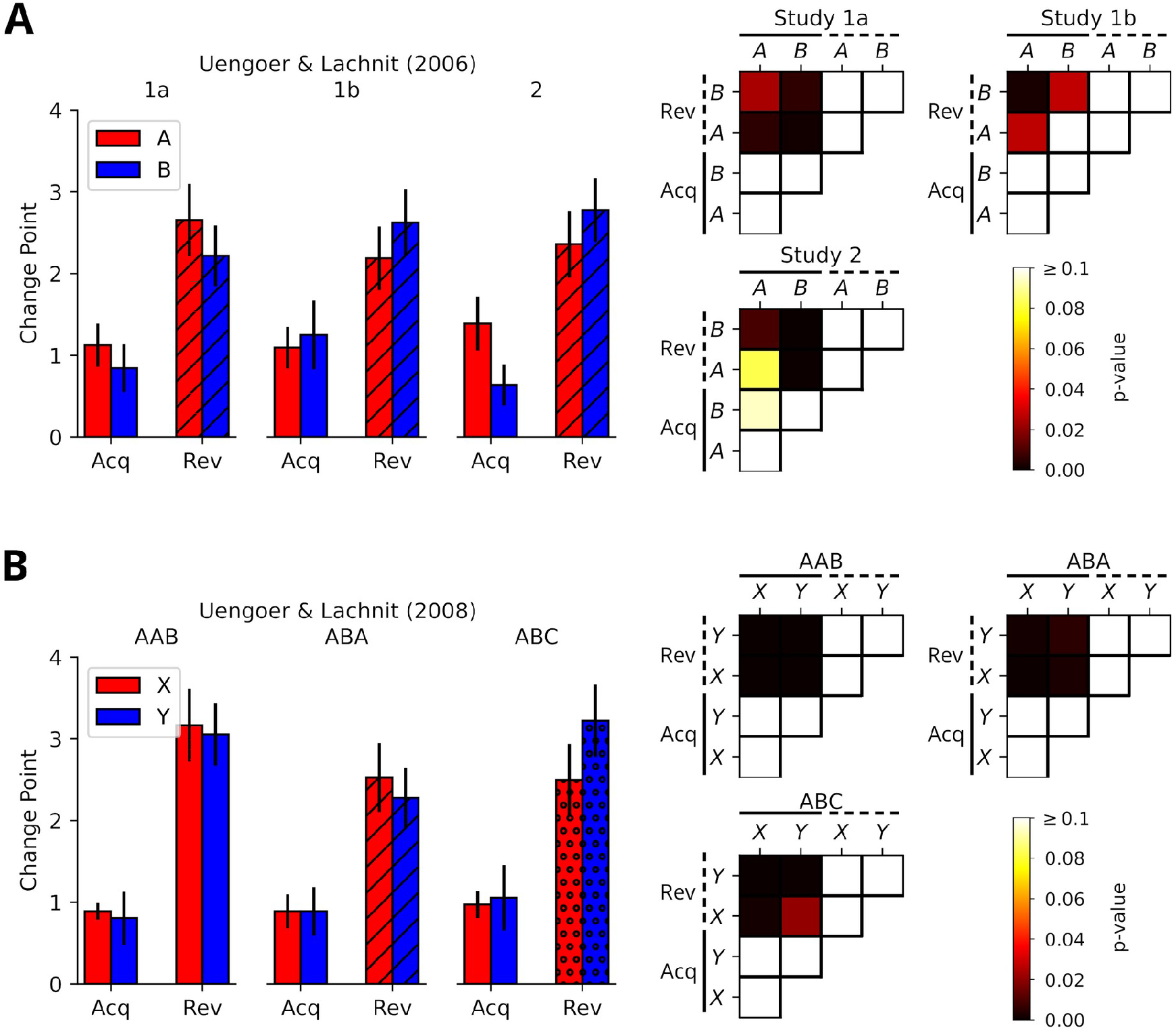
Detailed comparison of behavioral change points. **A**: Re-analysis of Uengoer and Lachnit (2006). Left: Shown are the average behavioral change points for the stimuli that underwent reversal in their experimental groups. Right: Pair-wise permutation tests of behavioral change points for the stimuli that underwent reversal for from acquisition (Acq, solid line) and reversal (Rev, dotted line) phases. **B**: Same as **A** but for Uengoer and Lachnit (2008).

**Figure S3:**
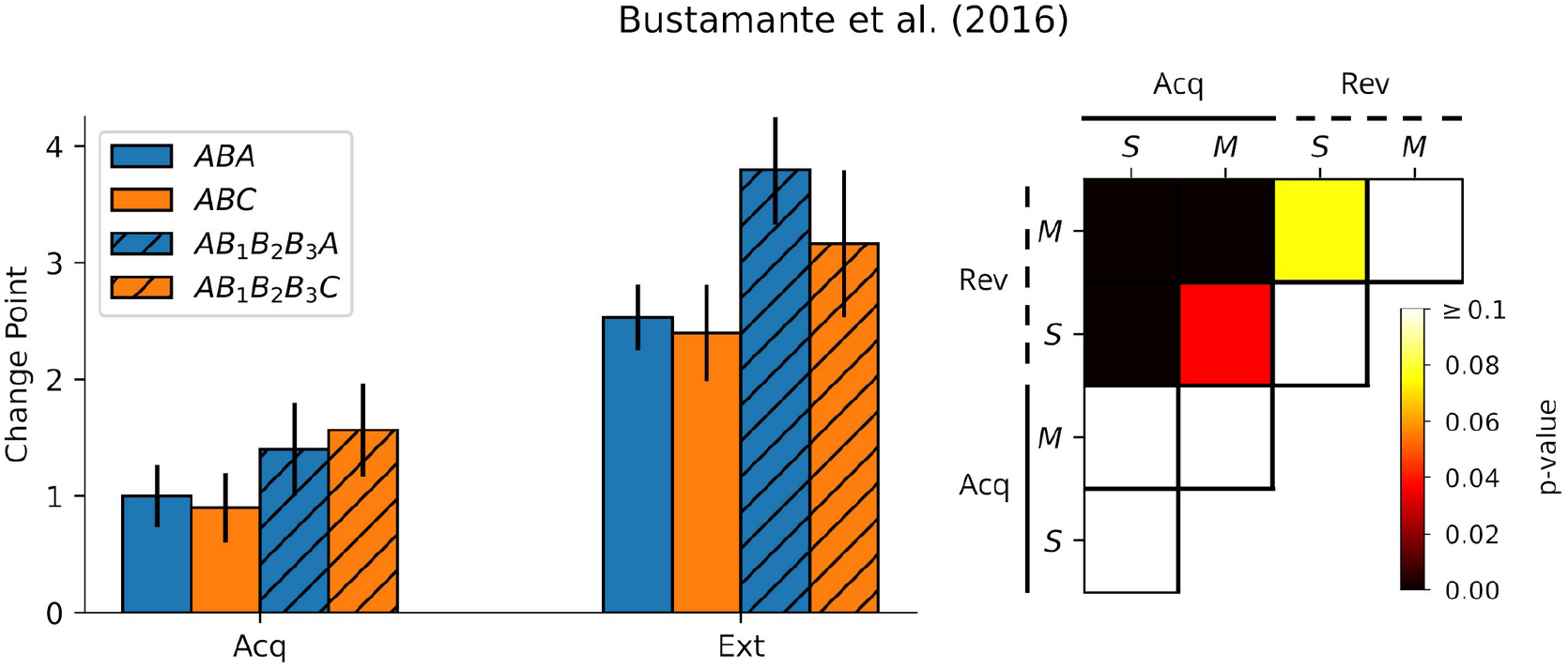
Using multiple contexts slows down extinction learning. **Left**: Average change points for the extinction stimulus during acquisition (Acq) and extinction (Ext) phases from Bustamante et al. (2016). Stripes indicate that extinction was performed using multiple contexts. **Right**: Heatmaps depicting the p-values obtained from permutation tests comparing change points for single-(S) and multiple-context (M) conditions in acquisition and extinction phases. P-values are corrected for multiple comparisons using the Holm-Bonferroni method.

**Figure S4:**
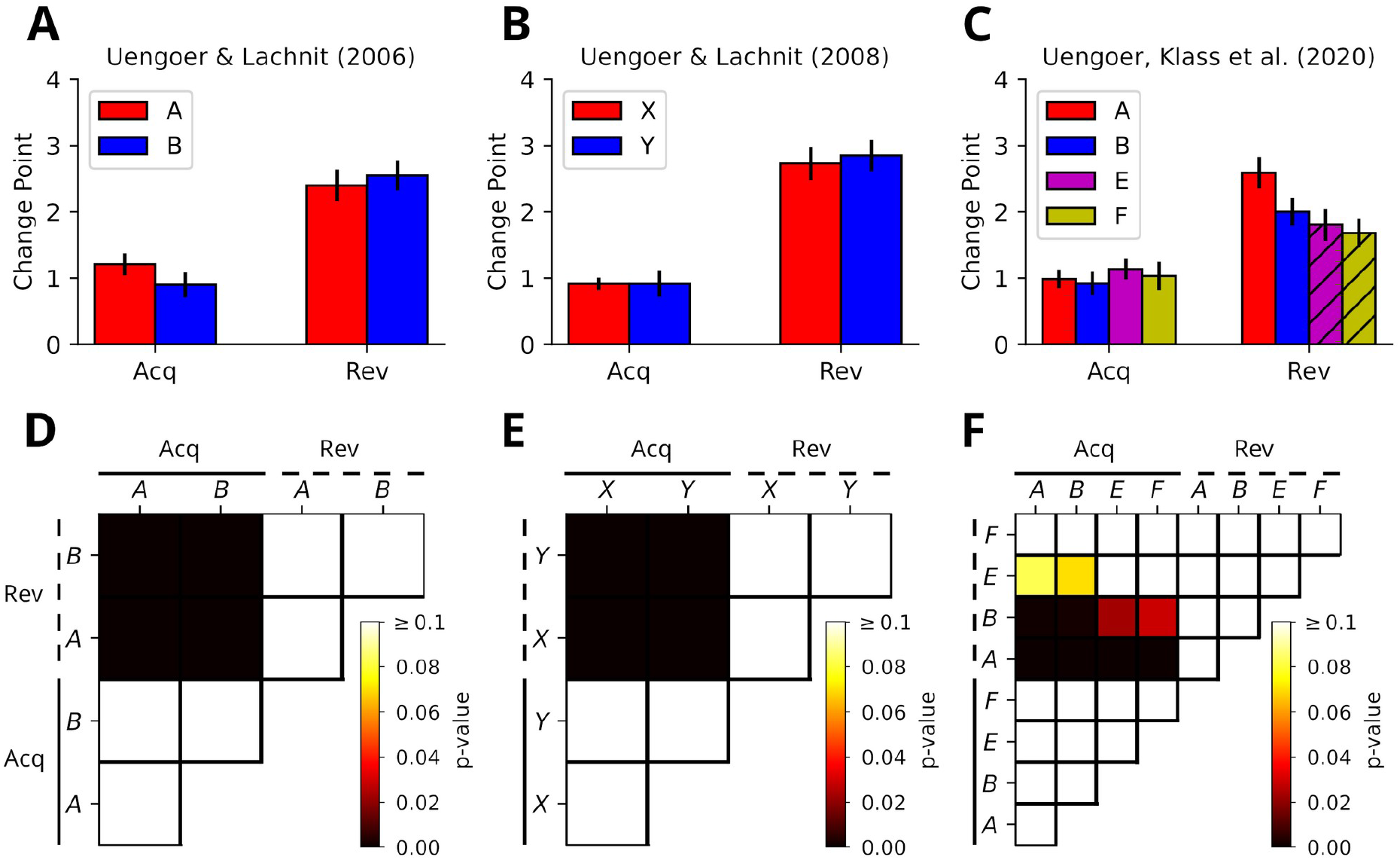
Reversal learning is slower compared to acquisition learning. **A** : The average change points for the reversal stimuli during acquisition (Acq) and reversal (Rev) in Üngör and Lachnit (2006). During acquisition the change points tend to occur around the first trial while during reversal they do so after two to three trials. **B**: Same as **A** but for Uengoer and Lachnit (2008). **C**: Same as **A** but for Uengoer et al (2020). Stripped bars indicate that reversal occurred in different context. **D-F**: Heatmaps depicting the p-values obtained from permutation tests for Üngör and Lachnit (2006, 2008); Uengoer et al. (2020), respectively. P-values are corrected for multiple comparisons using the Holm-Bonferroni method. Generally, change points significantly differ between phases, but not within phases.

**Figure S5:**
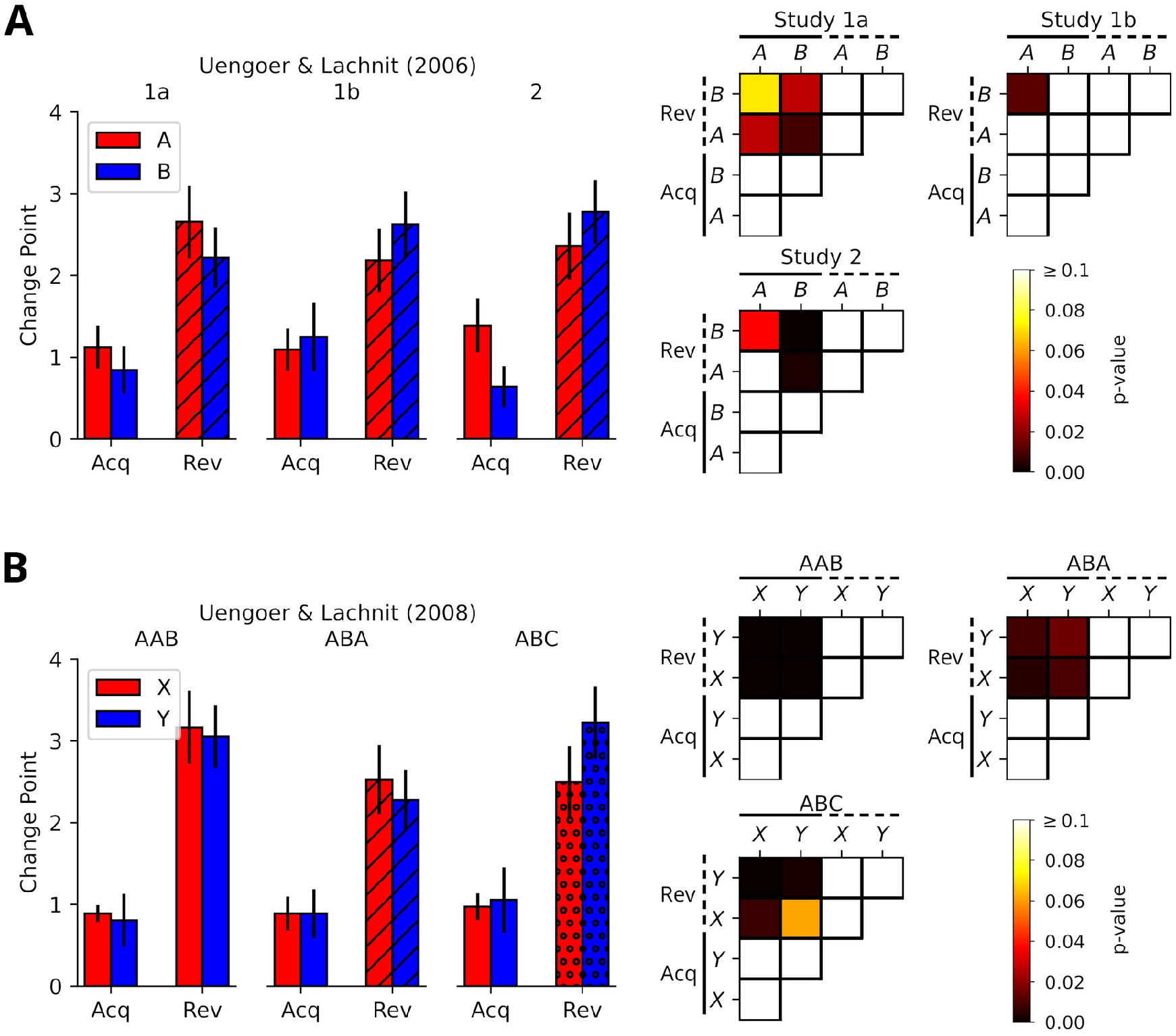
Detailed comparison of behavioral change points. **A**: Re-analysis of Uengoer and Lachnit (2006). Left: Shown are the average behavioral change points for the stimuli that underwent reversal in their experimental groups. Right: Pair-wise permutation tests of behavioral change points for the stimuli that underwent reversal for from acquisition (Acq, solid line) and reversal (Rev, dotted line) phases. P-values are corrected for multiple comparisons using the Holm-Bonferroni method. **B**: Same as **A** but for Uengoer and Lachnit (2008).

**Figure S6:**
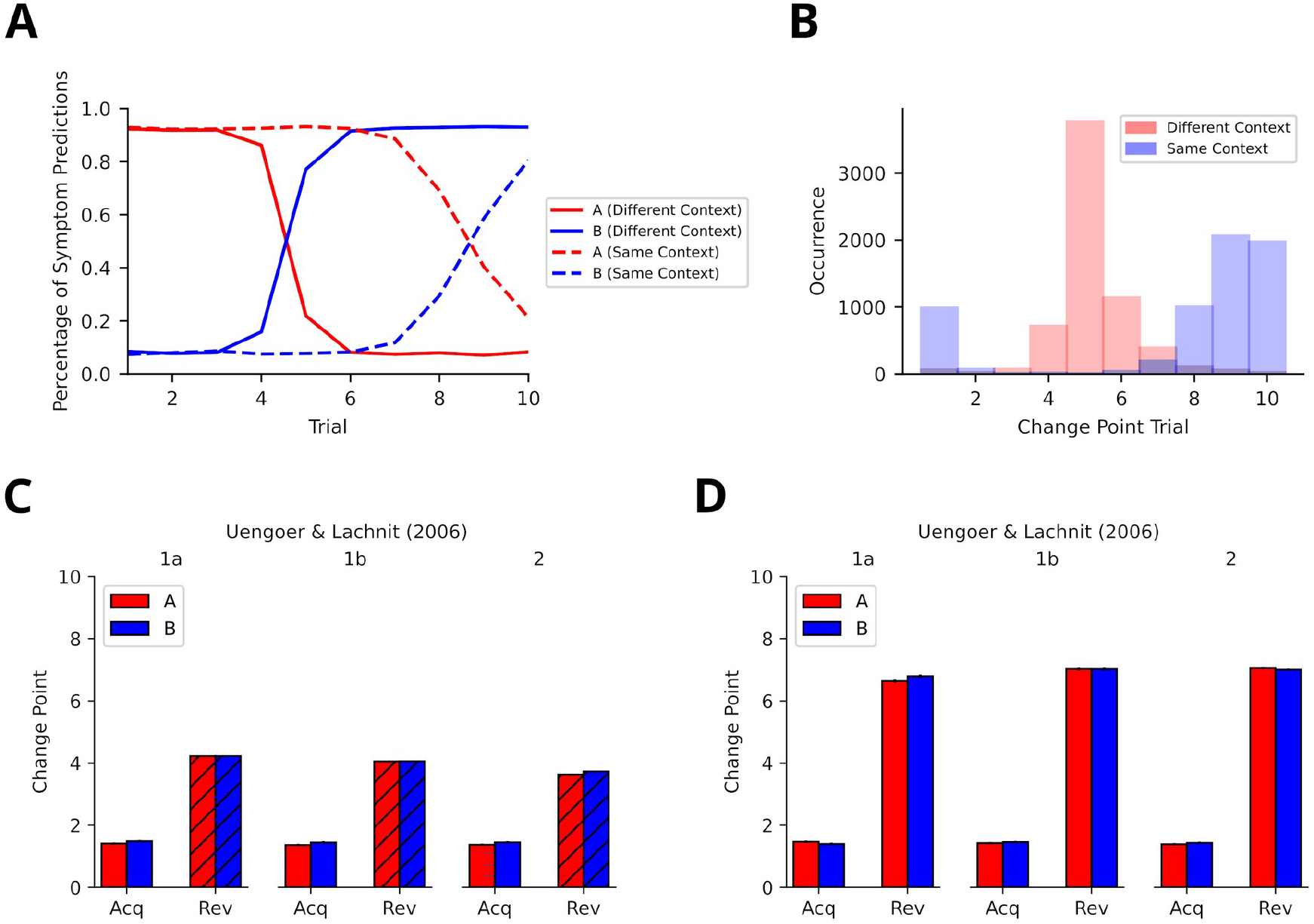
Replay of experiences slows down reversal learning when occurring in the same context. **A**: The average learning curves for stimuli A an B in the reversal phase for different and same context reversal learning simulations of study 1a from Uengoer and Lachnit (2006). A different parameter set was used since the fitted parameter yielded learning dynamics that were too fast to detect differences for reversal learning in the same vs a different context. Learning is considerably slower in the same context reversal learning simulation. B: The distributions of detected change points trials for different and same context reversal learning simulations of study 1a from Uengoer and Lachnit (2006). For the different context reversal learning simulation detected change points are clustered around trial five. For the same context reversal learning simulation change points are clustered around trials nine and ten. For some simulations reversal learning did not occur within ten trials which yielded some change points being detected at the first trial (recall that our analysis treats no change point as a change point at the first trial). **C:** The average change point trials for stimuli A and B in acquisition and reversal phases for simulations of the studies from Uengoer and Lachnit (2006). Change points shift towards later trials during the reversal phase. Striped bars indicate a change of context. **D:** Same as **C** but context did not change for the reversal phase. Change points consistently shift more during the reversal phase compared to **C**.

